# A new model for coordinating the functions of TIMELESS at the replication fork

**DOI:** 10.64898/2025.12.17.694947

**Authors:** Sameera Vipat, Rohan Harolikar, Karina Šapovalovaitė, Naga Raviteja Chavata, Syed Shahid Musvi, Arthur Morgunov, Sigvard Vällo, Tatiana N. Moiseeva

## Abstract

TIMELESS is an essential protein that supports a multitude of various cell functions, from replication fork progression through intrinsic barriers in the genome and DNA damage checkpoint to double strand break repair, transcription, and circadian rhythm. How TIMELESS coordinates its various roles at the replication fork and in DNA damage response, and how its canonical position at the leading edge of the replication fork could facilitate its role in DNA damage checkpoint, remain poorly understood. Using an auxin-inducible degron system, we show that TIMELESS-depleted cells exhibited S phase entry defects, and compromised chromatin loading of CLASPIN and TIPIN - the other two components of the Fork Protection Complex (FPC). We further show that FPC chromatin loading was concurrent with the activation of the replicative helicase, but also required proficient DNA synthesis. Proximity labelling experiments suggested the existence of more than one molecule of TIMELESS per replication fork. TIMELESS interaction with the replicative helicase was essential for the speed of replication fork progression, but not for the role of TIMELESS in the activation of the replication checkpoint. Our data propose a new model for coordinating essential functions of TIMELESS in replication fork progression and checkpoint activation.

**Graphical Abstract:** 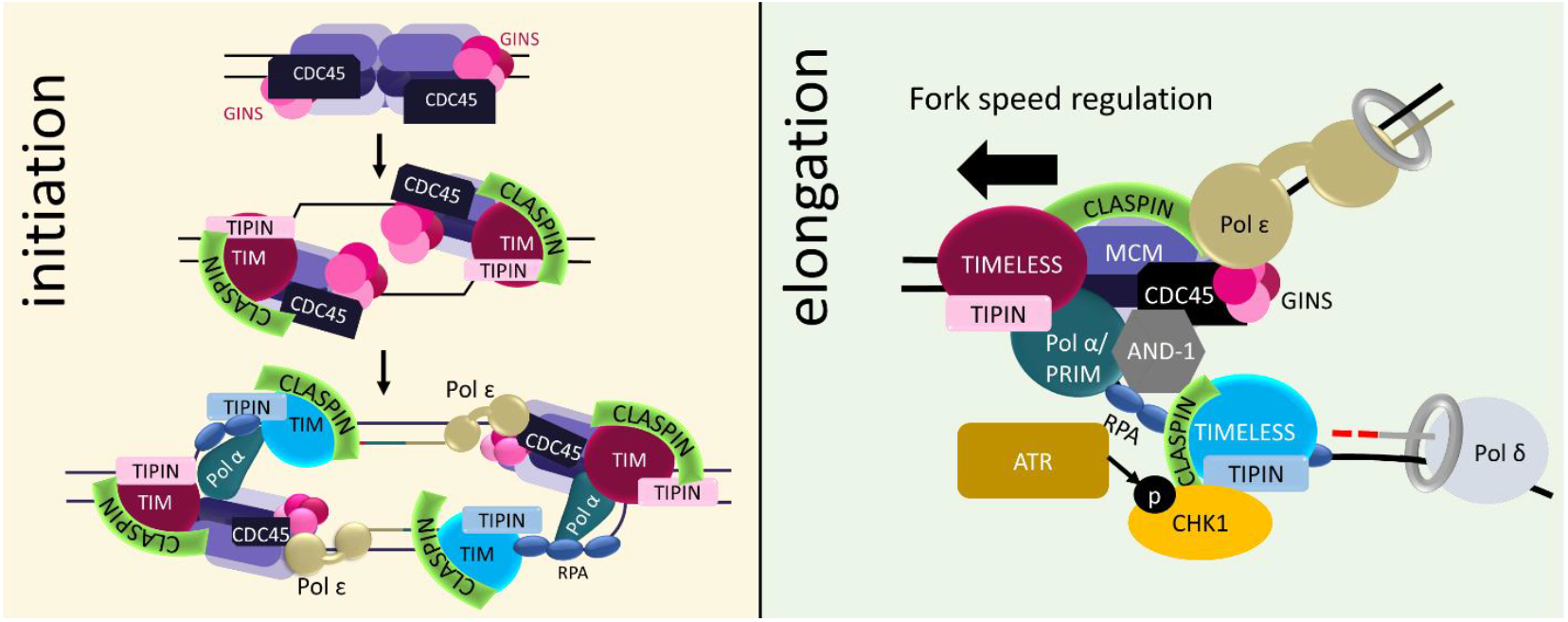

## Introduction

Timely and accurate DNA replication is essential for cell proliferation and genome stability. TIMELESS is a part of Fork Protection Complex (FPC, TIMELESS-TIPIN-CLASPIN) that ensures optimal replication fork speed and plays several functions in replication stress response (1–3), including replication checkpoint signaling (4) . TIMELESS performs most of its functions bound to its obligate binding partner TIPIN, and a depletion of one partner leads to a downregulation of the other (5). TIMELESS-TIPIN have been shown to coordinate the activities of the MCM helicase and the polymerases (6–9) by stimulating DNA polymerase activity and regulating MCM helicase activity at the same time (7, 10), ensuring the maintenance of normal replication fork speed.

Most mechanisms of replication initiation are conserved in eukaryotes (11, 12). The assembly of the replisome begins in the G1 phase of the cell cycle with origin licensing - loading inactive double hexamers of MCM helicase onto the origins marked by the Origin Recognition Complex (ORC). This is followed by CDK-dependent recruitment of CDC45 and GINS to MCM, leading to CMG helicase assembly. MCM is then activated by DDK- and CDK2-dependent phosphorylations and the action of MCM10 and RECQL4 (13). After RNA primers are synthesized by primase, DNA polymerase alpha (POLA) adds the first short stretch of DNA, after which PCNA is loaded by Replication Factor C (RFC) at the template-primer 3’ end sites (14). PCNA loading initiates DNA polymerase delta (POLD) loading (15), and POLD then takes over DNA synthesis (16). Polymerase epsilon (POLE) subsequently takes over the leading strand from POLD (17), and after this ‘polymerase switch’ step, two replication forks moving in opposite directions are established.

When the FPC is recruited to the replication complex is still poorly understood. One study indicated that TIMELESS is loaded to origins with the same dynamics as CDC45 (21), while another study reported that TIMELESS associates with MCM even before the beginning of S phase (1), which may indicate that it is loaded to origins along with MCM (18) in G1. The latter model is not consistent with the TIMELESS location at the leading edge of the fork which would be concealed in the MCM double hexamers according to structural data (21, 22). What role, if any, TIMELESS-TIPIN play in replication initiation is also not clear. Some studies showed excessive origin firing in TIMELESS-depleted cells (5) and others reported delayed S phase entry and a reduction in the S phase cell population (18) upon TIMELESS depletion. Here, using cell synchronizations and the mini-auxin inducible degron (mAID) system (19, 20), we investigate TIMELESS chromatin loading during replication initiation and its role in regulating DNA replication. We show that TIMELESS chromatin loading is initiated concurrently with CMG assembly in a CDK2/CDK1/CDC7-dependent manner, but is also partially dependent on DNA synthesis. We also show that TIMELESS depletion compromises chromatin loading of CLASPIN and TIPIN during the initiation of DNA replication resulting in a decrease in DNA synthesis in S phase.

There is conflicting evidence about the position of TIMELESS at the replication fork, with structural studies finding that TIMELESS is positioned ahead of the advancing MCM helicase (23–25), while biochemical evidence suggested that TIMELESS interacts with the polymerases at the fork and stimulates their activities (7, 10). Direct interaction between TIPIN and RPA/ssDNA (4) also suggests position for FPC at the ssDNA between Okazaki fragments. Here, we employ a split-TurboID-based proximity labelling system (26) to study the proteins in proximity to TIMELESS at the fork and better understand its position at the replisome. The set of proteins biotinylated by the used split-TurboID pairs supported the model in which there is an additional FPC at ssDNA on the lagging strand. Further, using a mutant of TIMELESS unable to interact with MCM but retaining its interactions with TIPIN and CLASPIN, we show that TIMELESS-MCM interaction is essential for the role of TIMELESS in replication fork speed, and for full chromatin loading of TIPIN and CLASPIN, but not essential for its role in checkpoint signaling.

TIMELESS overexpression is often detected in cancers, providing resistance to chemotherapies and stimulating epithelial to mesenchymal transition as well as metastases (3, 29–31); however, which roles of TIMELESS are important for cancer progression remains unclear. Our study provides insights into the coordination of various roles of TIMELESS at the replication fork, which may be important for separating its essential functions from the ones that could be potentially targeted by cancer therapies.

## Materials and methods

### Plasmids and cloning

For osTIR1 expression, we used pMK243-Tet-OsTIR1-PURO and AAVS1 CMV-OsTIR1(F74G) (a gift from Masato Kanemaki—Addgene plasmids # 72835 and #140536) plasmids (24, 51).

For mAID KI templates, homology arms were synthesized by Genscript in the pUC57 vector with the stop codon substituted with a BamHI site (5’ GGATCC 3’). The insert from plasmid pMK293 (mAID-mCherry2-Hygro) (Addgene plasmid # 72831 was a gift from Masato Kanemaki) (20), was cloned into the BamHI site of the synthesized plasmid, between the synthesized homology arms.

The TIMELESS C-terminus-targeting gRNA (ACGTGTCTATCTCTACCCCT) was expressed from the pSpCas9 BB-2A-Puro (PX459) v2.0 plasmid (Genscript).

TurboID fragments were cloned out of the 3xHA-TurboID-NLS_pCDNA3 plasmid that was a gift from Alice Ting (Addgene plasmid # 107171) (47) using the following primers TurboN_F: 5’-ATACGCGTTACCCCTATGACGTCCCAGA-3’; TurboN_R: 5’-ATGTTTAAACCAGAATCTGTTTAGCGTTCA-3’; TurboC_F: 5’-ATGATATCGGACAGCTGGACGGCGGGAG-3’; TurboC_R: 5’-ATGTTTAAACCTTTTCGGCAGACCGCAGAC-3’. TIMELESS ORF was cloned out of pcDNA4-Flag-Timeless, a gift from Aziz Sancar (Addgene plasmid # 22887) (48), using the following primers: Tim_Sgf1_F: 5’-ATGCGATCGCCATGGACTTGCACATGATGAA-3’ and Tim_Mlu1_R: 5’-ATACGCGTGTCATCCTCATCATCCTCAA-3’. AND-1 ORF was cloned out of the Origene plasmid #NM_007086 using SfaAI and MluI restriction sites. ORC6 ORF was cloned out of the Origene plasmid NM_014321 using SfaAI and MluI restriction sites.

For generation of the TIMELESS-M* mutant, the sequence corresponding to amino acids 276-302 was deleted from the TIMELESS gene, using Gibson Assembly-based cloning.

### Cell lines, cell culture, and transfections

U2OS (ATCC HTB-96) cells were grown in RPMI-1640 medium (Capricorn Scientific) supplemented with 10% FBS (GIBCO) and 1% penicillin-streptomycin (Invitrogen).

RPE-1 hTERT cells (ATCC CRL-4000) were grown in DMEM/F12 medium (GIBCO) supplemented with 10% FBS (GIBCO) and 1% penicillin-streptomycin (Invitrogen).

293T cells (ATCC CRL-3216) were grown in DMEM medium (GIBCO) supplemented with 10% FBS (GIBCO) and 1% penicillin-streptomycin (Invitrogen).

For generation of mAID1 TIMELESS-depleting cell lines by CRISPR, as per (23, 24, 54), U2OS-osTIR1 cells were transfected with gRNA and the mAID-mCherry-hygroR KI HR templates. The growth medium was changed 8 h after transfection, 2.5 µM DNAPK inhibitor was added for 48 h. KI cells were selected with hygromycin until the non-transfected control died, followed by single cell cloning and KI validation by PCR and western blot. Transfections were carried out using Lipofectamine 2000 (ThermoFisher), according to the manufacturer’s instructions. For generation of mAID2 TIMELESS-depleting cell lines by CRISPR, U2OS-based cells, expressing osTIR1 F74G mutant under the CMV promoter, were created by transfecting U2OS cells with the expression plasmid and selecting with puromycin, followed by the same mAID knock-in procedure.

For validations of the knock-ins, genomic DNA was isolated using genomic DNA miniprep kit (Zymo Research, D3025). Primers used to validate the presence of the knock-in were as follows: TIMhtestF: 5’-CGACAATTGCTGGACAGCGAC-3’ and gtestR: 5’-GGATCCTTACTTGTACAGCTC-3’; the primers used to validate the absence of unedited TIMELESS allele were as follows: TIMhtestF and TIMhtestR: 5’-ATCTCCAGAGAGCTGCTGGGG-3’.

For TIMELESS-M*-expressing cell line generation, mAID1 Clone 10 or Clone 28 cells were transfected with the corresponding plasmid using Lipofectamine 2000 (ThermoFisher), according to manufacturer’s instructions, growth medium was changed 8 h after transfection, and cells were selected with G418 until the non-transfected control died, followed by single-cell cloning and a validation by western blot.

### Cell lysis, insoluble chromatin isolation, and western blots

Cells were lysed in TGN lysis buffer (50 mM Tris-HCl (pH 7.5), 150 mM NaCl, 50 mM NaF, 1% Tween-20, 0.5% Nonidet P-40, and protease inhibitors (Pierce #A32953)) for 20 min on ice. Lysates were cleared by centrifugation, and soluble protein was used for immunoprecipitation or mixed with 2X Laemmli Sample Buffer (Bio-Rad) and incubated for 7 min at 96 °C, followed by western blot. For nuclease-insoluble chromatin, pellets were suspended in NIB buffer (150 mM HEPES (pH 7.9), 1.5 mM MgCl_2_, 10% glycerol, 150 mM potassium acetate, and protease inhibitors) containing 1 μl of universal nuclease for cell lysis (ThermoFisher, 88700) per 100 μl of the buffer and incubated for 10 min at 37 °C on a shaker. Nuclease-insoluble chromatin was pelleted by centrifugation, washed with water, and resuspended in Laemmli Sample Buffer.

For western blot analyses, proteins were separated in 8%, 10%, or 12% SDS-polyacrylamide gels in running buffer (25 mM Tris, 192 mM glycine, 0.1% SDS), transferred onto PVDF membrane (BioRad, #1620177) in transfer buffer (25 mM Tris, 192 mM glycine, 10% ethanol), blocked with 5% non-fat milk (BioRad, #1706404) in TBST (TBS ThermoFisher, #BP2471, 0.1% Tween-20), incubated with an appropriate dilution of the primary antibody overnight at 4 °C, washed with TBST buffer, incubated with secondary antibody for 1 h at room temperature, washed with TBST, and developed using SignalFire™ Elite ECL Reagent (Cell Signaling, #12757P) and ImageQuant LAS 4000 imager (GE Healthcare). Quantification of western blots was performed using Fiji/ImageJ (version 1.53u).

### Coimmunoprecipitation

293T cells were transfected with an empty vector or N-terminally FLAG-tagged wild-type TIMELESS and TIMELESS-M* expressing constructs, using Lipofectamine 2000 as per the manufacturer’s instructions. 48 h later, cells were lysed in TGN buffer as described above, and FLAG-tagged proteins were immunoprecipitated using anti-FLAG M2 affinity gel beads (Sigma-Aldrich), followed by elution with FLAG peptide (100 μg/ml in TGN buffer 2 h, 4°C).

### Split-Turbo ID experiments

293T cells were transfected using Lipofectamine2000, according to the instructions from the manufacturer. 48h after transfection, cells were incubated for 1h with 50 µM biotin, washed with PBS, and lysed in RIPA buffer (150 mM NaCl; 50 mM Tris-HCl pH 7.5; 1 % Triton X100; 0.1 % SDS, and protease inhibitors (Pierce #A32953)) on ice. After sonication using a Bioruptor (20 cycles, 30 seconds on; 30 seconds off), the lysates were cleared by centrifugation (14 000 g, 15 min) and incubated with streptavidin agarose (Thermofisher) for 16 h. The beads were then washed once with RIPA buffer, once with 1M NaCl, and two more times with RIPA buffer. Washed beads were then boiled with 1x Laemmli buffer (BioRad) and analyzed by western blot.

### Synchronizations and chromatin loading of replication proteins

In order to synchronize U2OS cells, 2 mM thymidine was added to ∼25% confluent cells for 24 h. After thymidine removal, cells were washed once with warm PBS and allowed to recover in fresh medium for 5 h. Nocodazole was then added for 12 h to arrest the cells in G2/M. Dox (2 µg/ml) and 3-IAA (500 μM) for mAID1, or 5-pH-IAA (1.25 μM) for mAID2, were added at the same time as nocodazole. After release from nocodazole, cells were washed once with warm PBS, and incubated in pre-warmed medium with dox/aux or 5-Ph-IAA for the indicated periods of time. For the experiments with added inhibitors, cells were released from nocodazole, washed once with warm PBS, and incubated in pre-warmed medium with 5 μM of each inhibitor for the indicated periods of time.

For RPE-hTERT synchronizations, cells were grown to confluency, growth medium was replaced with one without FBS and cells were incubated for 48 h to achieve G0 arrest. To release from G0, cells were passed 1:5 into complete medium. Cells entered S-phase between 15 and 18 h after release from G0.

In order to assess the loading of the various proteins on chromatin, samples were collected by trypsinization at the indicated time points, pellets washed once with ice-cold PBS and kept at −80 °C. To obtain the nuclease-insoluble fraction, thawed pellets were resuspended in CSK buffer (10 mM PIPES pH 7.0, 300 mM sucrose, 100 mM NaCl, 3 mM MgCl_2_, 0.5 % Triton X-100, and protease inhibitors (Pierce #A32953)), incubated for 5 min on ice, followed by a 5-min centrifugation (1000 × g, 4 °C). pellets were washed once more with CSK buffer, digested in CSK buffer with universal nuclease for cell lysis (ThermoFisher, #88700) for 10 min at 37 °C. Samples were mixed with 2x Laemmli Sample Buffer (BioRad) and boiled for 10 min before proceeding to western blot analysis. Quantifications were performed using ImageJ and GraphPad Prism 9.

### Antibodies

POLE1 (Santa Cruz, #sc-390785, 1:500), GAPDH (Santa Cruz, #sc-47724, 1:1000), pCHK1 (Cell Signaling, #2360S, 1:1000), MCM4 (Cell Signaling, #3228S, 1:300), CDC45 (Santa Cruz, #sc-55569, 1:500), SLD5 (Santa Cruz, #sc-398784, 1:300), H3 (Santa Cruz, #sc-517576, 1:1000), POLE2 (Santa Cruz, #sc-398582, 1:500), PCNA (Santa Cruz, #sc-56, 1:1000), FLAG (Sigma, F3165-1MG, 1:3000), polD1 (Santa Cruz, # sc-374025, 1:500), TIMELESS (Santa Cruz, #sc-393122), TIPIN (Santa Cruz, #sc-135580), CLASPIN (Santa Cruz, #sc-376773), H2AX (Santa Cruz, #sc-517336), MCM2 (Santa Cruz, #sc-373702), MCM7 (Santa Cruz, #sc-9966), RPA32 (Santa Cruz, #sc-56770), POLA1 (Santa Cruz, #sc-373884).

### Inhibitors

ATRi AZD6738 (AstraZeneca), 5μM; DNAPKi NU7026 (Selleckchem), 2.5 μM; CDK1i Ro 3306 (Selleckchem), 5μM; CDK2i CVT-313 (Selleckchem), 5 μM; CDC7i XL413 (Selleckchem), 5μM; aphidicolin (MilliporeSigma), 2μM.

### Flow cytometry

For EdU FACS, cells were treated with 10 μM EdU for 30 min, trypsinized, washed with PBS, and fixed with cold 70% ethanol on ice for 30 min or at -20°C overnight. Cells were washed with PBS, and EdU staining was performed by using the EdU Click-iT kit (ThermoFisher, #C10632), according to the manufacturer’s instructions. For DNA staining 7-AAD (7-Aminoactinomycin D) (ThermoFisher, # A1310), FxCycle Far Red Stain (ThermoFisher, #F10348) or FxCycle™ PI/RNase Staining Solution (ThermoFisher, #F10797) were used. Samples were analyzed on FACSCalibur flow cytometer, and data were analyzed by using FCSalyzer software. Alternatively, the samples were analyzed on Cytoflex 2L followed by data analyzes using FlowJo. Software. GraphPad Prism 9 was used for statistical analyses.

### DNA fiber analysis

Cells were pulsed with 20 μM CldU followed by 200 μM IdU to mark ongoing DNA synthesis. After trypsinization and PBS wash, cells were lysed by adding 6 μl of lysis buffer (200 mM Tris–HCl pH 7.4, 50 mM EDTA, 0.5% SDS) to 2ul of cell suspension and spread carefully with a plastic tip on a glass slide. After a 5 min incubation, the DNA was allowed to slide down the tilted slide. After drying the slides for 10 min at room temperature, the DNA was fixed by incubation in 3:1 methanol-acetic acid for 5 min, and dried for 8 minutes at room temperature. DNA was then rehydrated in PBS during 2 5-min incubations. DNA was then denatured in 2.5N HCl for 1h, followed by 2 5-min PBS washes. The samples were blocked in 5% BSA-0.1% Triton X-100 for 1h at 37°C, and incubated with primary antibodies (CldU #ab6326 1:50 and IdU BD#3475801:66.6) at 4°C overnight. Slides were washed 4 times 5 min with PBS-0.1% Tween20 and incubated with secondary antibodies (1:150) for 1h at 37°C, washed 4 times 5 min with PBS-0.1% Tween20 and 2 times 5 min with PBS. The samples were then mounted with ProLong Diamond Antifade Mountant (ThermoFisher #P36961). The fibers were imaged with Nikon fluorescent microscope, and analyzed using ImageJ, at least 100 fibers per condition were measured.

## Results

### TIMELESS depletion slows down DNA synthesis during S-phase entry

In order to study the role of TIMELESS in replication initiation in human cells, we used a mini-auxin-inducible degron (mAID) system to achieve rapid and near-complete depletion of TIMELESS. In this system, the C-terminus of the target protein TIMELESS is tagged with a mAID-mCherry tag using CRISPR genome modification, and additionally the F-box protein osTIR1 is stably transfected into the cells (20). Original mAID system (mAID1) uses doxycycline-inducible wild-type osTIR1 (20) while improved version 2 (mAID2) uses constitutively expressed osTIR1 F74G allowing for a faster degradation (16 hours for mAID1 and 3 hours for mAID2) by eliminating the need to induce osTIR1 expression (19).

Two U2OS-based clones for mAID1-Clone 10 and Clone 28, and two clones for mAID2 - Clone O3 and Clone L4 were selected, based on their ability to deplete TIMELESS to near-complete levels, and, as expected, its partner TIPIN was concurrently downregulated (**Fig. 1A, B**). The levels of CLASPIN, the other FPC component, were not affected by TIMELESS depletion. mAID tagging did not affect the cell cycle or (**Fig. S1A**).

**Figure 1.**
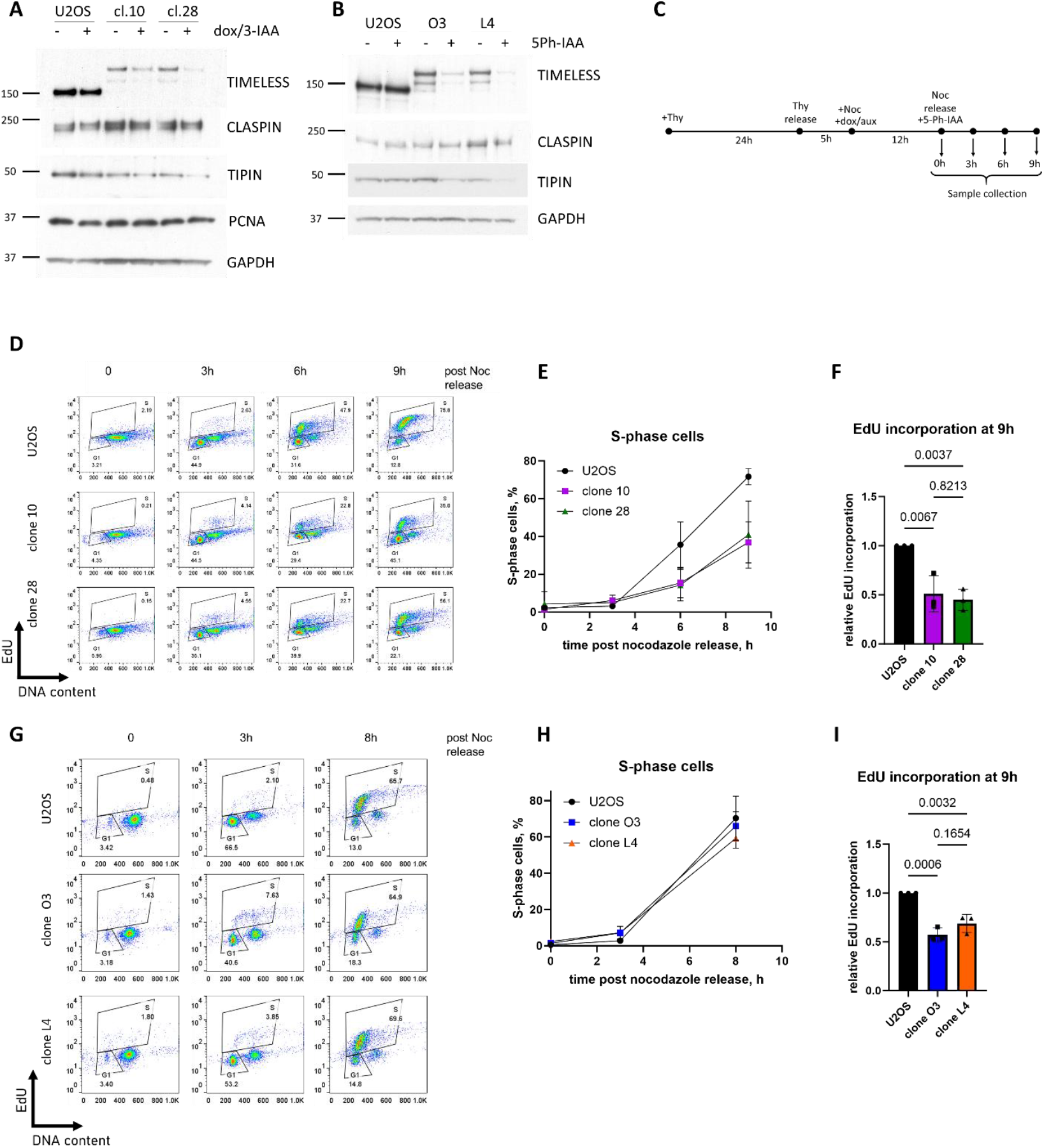
Effect of TIMELESS depletion on S-phase entry. **A-B**. U2OS, mAID1 clones Clone 10, Clone 28 or mAID2 clones Clone O3, Clone L4 were treated for 16 h with doxycycline and auxin 3-IAA, or with auxin 5-pH-IAA. Western blots of total cell lysates are shown (**A**,**B**). Synchronization timeline using the double thymidine-nocodazole block and the timepoints used for mAID1 (dox/aux) and mAID2 (5-Ph-IAA) clones (**C**). **D-I**. U2OS, mAID1 clones 10 and 28 or mAID2 clones O3 and L4 were synchronized using the double thymidine/nocodazole block, and treated as indicated on **C**. 10 µM EdU was added for the last 30 min of treatment. Flow cytometry plots showing EdU incorporation and DNA content (7-AAD staining) are shown (**D**,**G**). Quantification of the flow cytometry data is shown-mean + SD from n = 3 independent experiments (**E**,**F**,**H**,**I**), one-way ANOVA was used for statistical analyses.

TIMELESS depletion significantly affected the growth of both mAID1 and mAID2 clones, in agreement with the essential role of TIMELESS in cell proliferation (**Fig. S1B**). Doxycycline treatment slightly reduced the growth of U2OS cells in the mAID1 experiment, as has been previously observed (27, 28), while 5-Ph-IAA did not have any effect on U2OS in the mAID2 experiment (**Fig. S1c**).

Next, in order to investigate the effect of TIMELESS depletion on S phase entry, we depleted TIMELESS in cells that were synchronized in G2/M using a double thymidine-nocodazole block and then allowed to enter the S phase at the same time (**Fig. 1C**). We added dox/3-IAA to mAID1 cells at the time of nocodazole addition and 5-Ph-IAA to mAID2 cells at the time of release from nocodazole. Cells were collected at the indicated timepoints following the release from nocodazole, marking G2/M, G2, and S-phase entry. mAID tagging had no effect on the cell cycle progression after synchronization without TIMELESS depletion (**Fig. S1D**). Quantifications of S phase cells (**Fig. 1E, H**) and G1 phase cells (**Fig. S1E-F**) showed a significant S-phase delay in mAID1 clones 10 and 28, but not in mAID2 clones O3 and L4 after TIMELESS depletion. Additionally, DNA synthesis as measured by the level of EdU incorporation was lower in all TIMELESS-depleted clones compared to U2OS cells (**Fig. 1F, I**), which may be at least partly due to the previously documented (29) slower fork speed upon TIMELESS depletion.

The unexpected difference between mAID1 and mAID2 clones led us to consider possible differences between these systems. mAID1 clones needed to be incubated with 2ug/ml doxycycline to induce osTIR1 expression, and these cells were therefore depleted of TIMELESS during M/G1 transition, while mAID2 clones only depleted TIMELESS in late G1. As origin licensing happens during early G1 phase, we tested whether licensing is affected by TIMELESS depletion in all four clones (**Fig. S1G**). In this experiment cells were incubated with dox/3-IAA (mIAD1) or 5-Ph-IAA (mAID2) for 16h, leaving mAID2 clones without TIMELESS for a longer period of time than mAID1 clones. Despite this, we observed a clear delay in licensing specifically in clones 10 and 28 (mAID1 clones), but not in mAID2 clones O3 and L4. This result led us to hypothesize that it is not TIMELESS depletion that affected origin licensing, but perhaps a doxycycline treatment. To confirm this, we treated mAID2 clone O3 with doxycycline – alone or in combination with 5-Ph-IAA, and observed MCM chromatin loading defect in response to 16 h of 2 ug/ml doxycycline treatment, independently of TIMELESS depletion (**Fig. S1H**). These experiments suggest that doxycycline treatment in mAID1 system delays origin licensing resulting in S-phase delay.

### TIMELESS depletion leads to a defect in chromatin loading of CLASPIN

We next aimed to determine whether TIMELESS depletion affected chromatin loading of any replisome components. We synchronized mAID clones and U2OS cells using the double thymidine-nocodazole block, depleted TIMELESS as described above, and analyzed the presence of selected replication proteins in the chromatin fraction using western blot at indicated timepoints. As expected, low levels of TIMELESS and TIPIN resulted in their absence on chromatin in S-phase cells as well. Interestingly, we also observed a delay in the chromatin recruitment of CLASPIN (in all TIMELESS-depleted clones) and PCNA (in mAID1 clones only) (**Fig. 2A-D**) during S-phase entry. CDC45 chromatin loading during S-phase entry was not affected by TIMELESS depletion, indicating that there was no noticeable delay in CMG assembly. Our data showed that TIMELESS depletion affects the chromatin loading of CLASPIN during S-phase entry.

**Figure 2.**
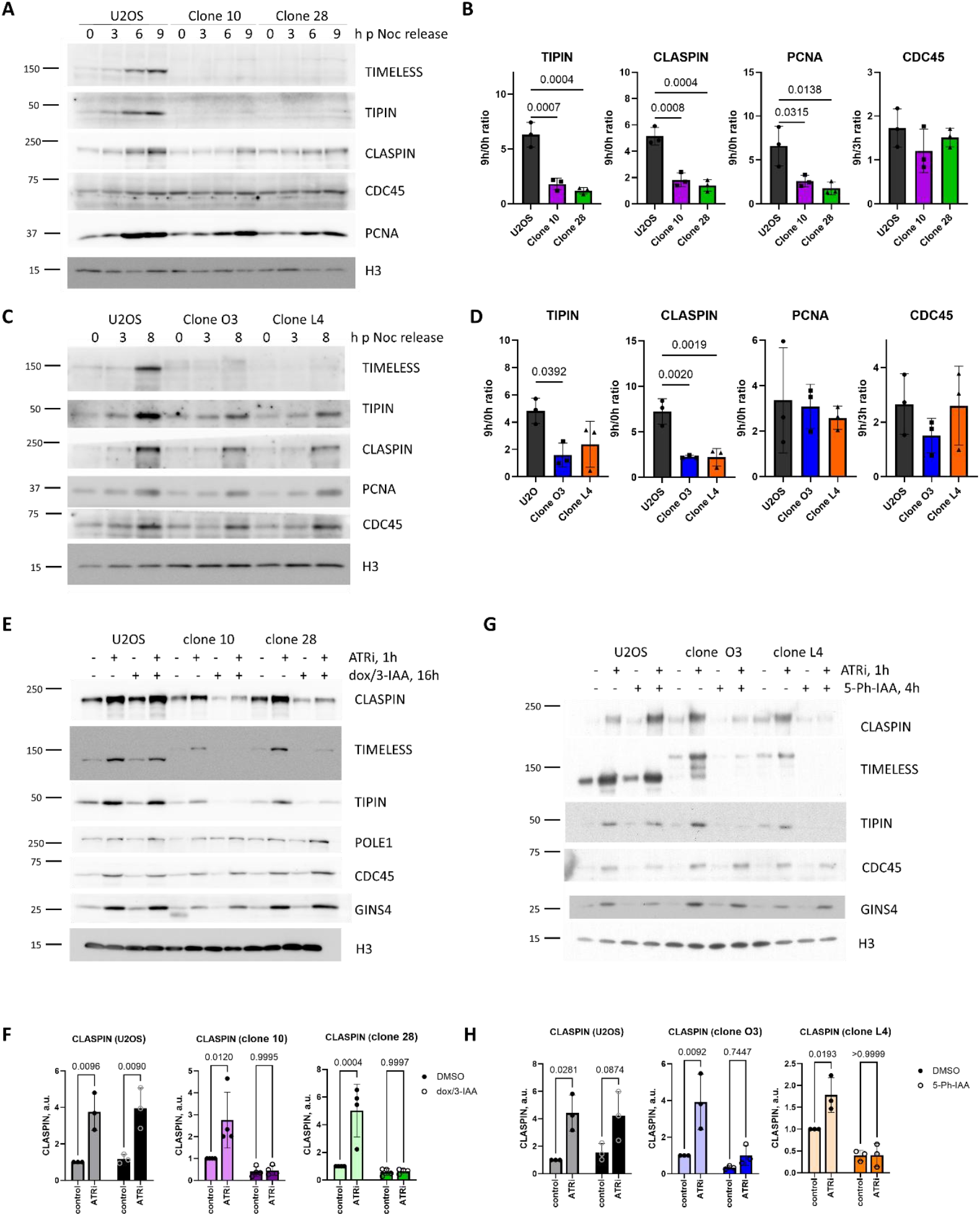
Effect of TIMELESS depletion on FPC chromatin loading during replication initiation. **A-D**. U2OS, mAID1 clones 10 and 28, or mAID2 clones O3 and L4 were synchronized and treated as shown on **Fig. 1C**. Western blot analysis of the nuclease-insoluble chromatin fraction from the cells collected at the indicated timepoints is shown (**A**,**B**). Quantifications of protein loading are shown (**C**,**D**). Mean + SD from 3 independent experiments is shown. **E-H**. U2OS, mAID1 clones 10 and 28, or mAID2 clones O3 and L4 were treated for 16h with dox/3-IAA, or for 4h with 5-Ph-IAA, as indicated. 5 µM ATRi was added to the indicated samples for 1 h, followed by cell lysis and the isolation of the nuclease-insoluble chromatin fraction. Western blots of the insoluble chromatin fraction are shown (**E**,**G**). Quantifications of protein loading from E,F are shown (**F**,**H**). Mean + SD from 3-4 independent experiments is shown, one-way ANOVA (**B**,**D**) or two-way ANOVA (**F, H**) was used for statistical analyses.

To confirm our findings using an alternative approach, we used an ATR inhibitor AZD6738 (ATRi) which is known to rapidly induce massive origin firing resulting in a strong recruitment of replication proteins to chromatin, MCM4 phosphorylation and an increase in EdU incorporation by replicating cells (35). We depleted TIMELESS in mAID clones and added DMSO or 5 µM ATRi (AZD6738) for the last 1 h to induce origin firing. TIMELESS depletion slightly dampened but did not prevent the increase in EdU incorporation in response to ATRi (**Fig S2A, B**). We observed significant reductions in CLASPIN and TIPIN recruitment to chromatin in response to ATRi in both sets of mAID clones (**Fig. 2E-H**), while the recruitment of CDC45 and GINS remained unaffected.

These data demonstrate that TIMELESS was required for the recruitment of CLASPIN to chromatin during the initiation of DNA replication for the proper assembly of the FPC at the replication fork.

### TIMELESS chromatin loading during origin firing requires CMG assembly and the initiation of DNA synthesis

To gain more insight into FPC recruitment to replication complexes during S-phase entry, we studied the chromatin loading of TIMELESS, TIPIN and CLASPIN relative to the loading of other replication proteins using cell synchronization.

We synchronized U2OS using the double thymidine-nocodazole block, collected cells at the indicated timepoints after their release from nocodazole (from G2 through G1 and S-phase entry), and analyzed the chromatin fraction by western blot (**Fig. 3A,B)**. TIMELESS and CLASPIN loaded concurrently with CDC45 (a marker of CMG assembly), but notably before DNA polymerase alpha, indicating that DNA synthesis was not required to initiate FPC chromatin loading during S-phase entry. Similar results were obtained with RPE1-hTERT cells synchronized using serum starvation and contact inhibition, confirming that TIMELESS chromatin loading was initiated before DNA polymerase alpha recruitment (**Fig. S3C-D**).

**Figure 3.**
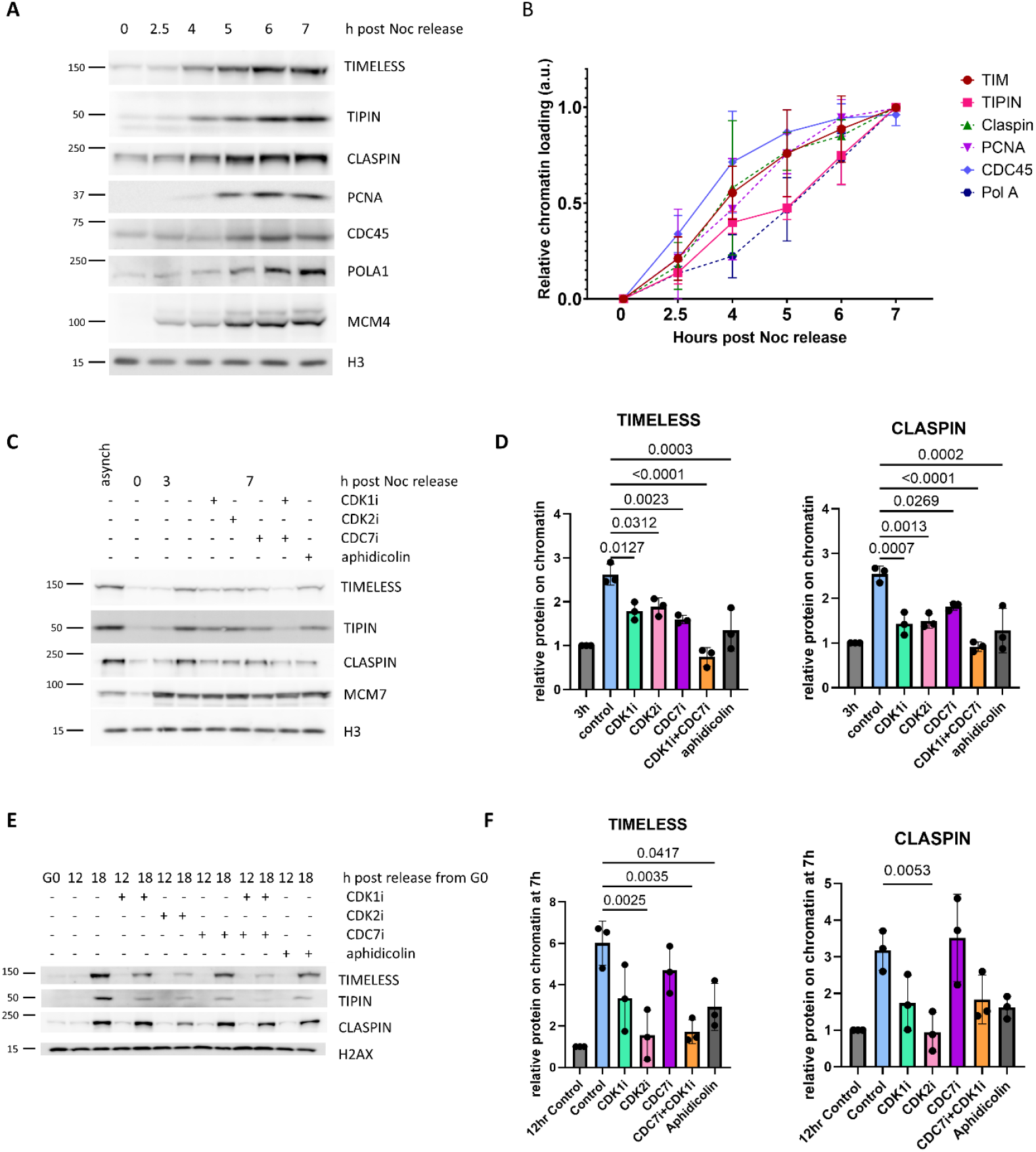
Timing and regulation of FPC chromatin loading during replication initiation. **A-D**. U2OS cells were synchronized as indicated on **Fig. 1C**. Samples were collected at the indicated timepoints post nocodazole release. Western blot analysis of the chromatin fraction from the cells collected at the indicated timepoints is shown (**A)**. Quantifications of protein loading from A are shown -mean + SD from 3 independent experiments (**B**). **C-D**. 3h post nocodazole release, the indicated inhibitors were added. Samples were collected at the indicated timepoints post nocodazole release. Western blot analysis of the nuclease-insoluble chromatin fraction from the cells is shown (**C**). Equal amounts of protein were loaded. Quantifications of protein loading from C are shown as mean + SD from 3 independent experiments **(D). E**. RPE-hTERT cells were synchronized by contact inhibition and serum starvation as indicated on **Fig S3C**. Indicated inhibitors were added 10h after release from G0, samples were collected at indicated timepoints, western blot analyses on chromatin fraction are shown (**E**). Quantifications of protein loading from **E** are shown -mean + SD from 3 independent experiments (**F**). One-way ANOVA was used for statistical analyses.

Further, to identify the steps of replication initiation required for FPC chromatin loading, we treated U2OS cells with CDK1, CDK2 or CDC7 inhibitors, or the B-family DNA polymerase inhibitor aphidicolin. Cell cycle analysis confirmed that the inhibitors we used affected S phase entry (**Fig. S3A-B**). Similar results were obtained in non-cancerous RPE1-hTERT cells (**Fig. S3E**). FPC chromatin loading was affected by CDK1, CDK2 and CDC7 inhibitors, and the combination of CDK1i and CDC7i almost completely suppressed FPC chromatin loading (**Fig. 3C-F**) in both cell lines. Since these kinase activities are known to regulate CMG assembly[28], we concluded that CMG assembly was likely required for FPC chromatin loading (**Fig. 3C-F**). Interestingly, aphidicolin also decreased FPC chromatin loading, indicating that DNA synthesis was needed for the stable association of FPC with chromatin (**Fig. 3C-F**). Given that TIMELESS and CLASPIN associated with chromatin before POLA was recruited (**Fig 3A, B**), DNA synthesis could not be the step that initiated FPC loading. Our data indicate that there are two steps of replication initiation that regulate FPC association with chromatin – CMG assembly and the initiation of DNA synthesis.

### Proximity labelling enabled by TIMELESS interaction with core replication initiation proteins indicates the presence of more than one molecule of TIMELESS per fork

In order to shed some light on the position of TIMELESS in the replisome and possible mechanisms of recruitment, we decided to look into the proteins located in proximity to TIMELESS at the replication fork. We therefore employed a split-Turbo ID-based proximity labelling strategy (26), allowing biotinylation of nearby proteins only when two bait proteins tagged with two complementary parts of TurboID biotin ligase (TurboN and TurboC) come together to form a functional enzyme. To identify suitable replication proteins to be combined with TIMELESS for this approach, we tagged several replication proteins (MCM4, AND-1, ORC6, POLE2) with TurboC and combined each of them with TurboN-TIMELESS. TIMELESS-ORC6 and TIMELESS-AND-1 combinations yielded the best overall biotinylation signal (**Fig. 4A, S4A**). AND-1 is a trimer positioned on the other side of the CMG helicase with respect to the N-terminus of TIMELESS, making it an unlikely successful combination. ORC6 is a subunit of the Origin Recognition Complex, which does not travel with the replication fork. However, ORC6 has been shown to also participate in mismatch repair behind the fork (24), and if this ORC6 molecule forms a successful split-TurboID pair with TIMELESS, TIMELESS must be positioned close to one of the daughter strands.

**Figure 4.**
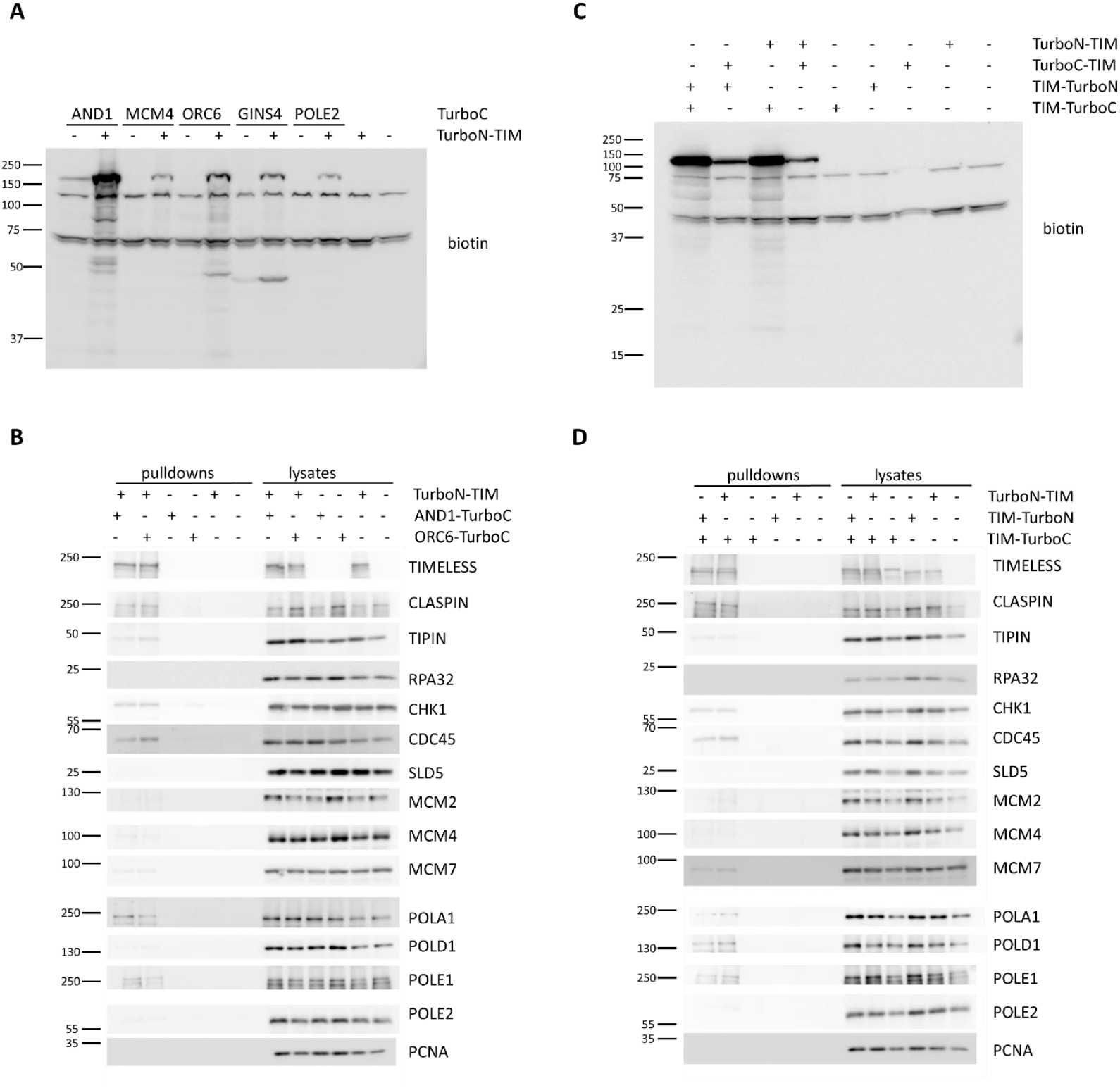
Split-TurboID approach uncovers TIMELESS-proximal proteins at the replication fork. **A-D**. HEK293T cells were transfected with the plasmids expressing indicated TurboN- or TurboC-tagged proteins. 48h after transfection cells were treated with 50 µM biotin for 1 hour and lysed. (**A, C**) Cell lysates were analyzed by western blot using antibodies against biotin (**A, C**). Western blot analyses of the streptavidin pulldowns and the lysates’ samples are shown (**B, D**).

To clarify the locations of contacts for the two successful combinations, we performed a streptavidin pulldown, and looked for known components of the replication fork among the proteins biotinylated by TIMELESS-AND-1 and TIMELESS-ORC6 pairs. Apart from the FPC components, the proteins showing the strongest biotinylation were CDC45 and CHK1, followed by DNA polymerases alpha and epsilon (**Fig. 4B**). Interestingly, we did not observe any noticeable differences in biotinylation levels for the selected proteins between the two combinations used in the experiment. Biotinylation of DNA polymerase epsilon on the leading strand and CHK1 (likely located close to ssDNA on the lagging strand) was difficult to reconcile with the canonical position of TIMELESS at the leading edge of the replication fork on the parental DNA.

While this canonical position of TIMELESS has been repeatedly demonstrated beyond any doubt (24, 25, 32), we decided to check if there could possibly be more than one TIMELESS molecule per replication fork.

To this end, we used split-TurboID combinations in which two TurboID parts were fused to two molecules of TIMELESS, both N-terminal and C-terminal tagging were tested (**Fig. 4C, S4B**). Interestingly, TIMELESS-TurboC produced strong biotinylation in combination with either TurboN-TIMELESS or TIMELESS-TurboN, indicating two molecules of TIMELESS being positioned in proximity to each other. TurboC-TIMELESS did not produce strong biotinylation in any combinations, probably due to TurboC being a larger tag that could be disruptive at the N-terminus of TIMELESS.

Analysis of proteins biotinylated by TIMELESS-TIMELESS combinations confirmed the presence of the same set of proteins that we saw biotinylated by TIMELESS-AND-1 and TIMELESS-ORC6 combinations (**Fig. 4D**). Interestingly, in addition to these proteins, the TIMELESS-TIMELESS combinations also biotinylated DNA polymerase delta, placing at least one molecule of TIMELESS in proximity to the lagging strand. Given that TIPIN was shown to interact with RPA and ssDNA (4), we propose that in addition to the leading edge FPCs, there could be secondary TIMELESS/TIPIN complexes at the ssDNA gaps on the lagging strand.

### Interaction between MCM and TIMELESS is essential for the full speed of replication forks, but not for the checkpoint signaling

In our proposed model at least two TIMELESS/TIPIN complexes are present at the replication fork, however, only one molecule of TIMELESS interacts with MCM. To clarify whether TIMELESS-MCM interaction is needed for the roles of TIMELESS in DNA replication, we generated a mutant of TIMELESS unable to interact with MCM - TIMELESS-M*. We designed it based on the data from (7), showing that TIMELESS amino acids 1-663 were sufficient for MCM interaction, but neither of the two broader regions present on either side of region 1-663 (1-307 or 308-1208) was sufficient for MCM interaction (7). We deleted the amino acids 275-301 forming a loop next to the MCM hexamer (**Fig. 5A**). We also took into consideration structural data (33), (32), and used Phyre2 (34) and AlphaFold (35) to ensure our deletion would cause minimal changes to the 3D structure of TIMELESS, aiming to preserve its interactions with its other partners. This mutant was predicted to have weaker interactions with MCM using Alphafold3 (**Fig. S5A**), and the disruption of TIMELESS-M* interaction with MCM was further confirmed using co-immunoprecipitation, while its interactions with TIPIN, CLASPIN and POLE were not affected (**Fig. 5B**).

**Figure 5.**
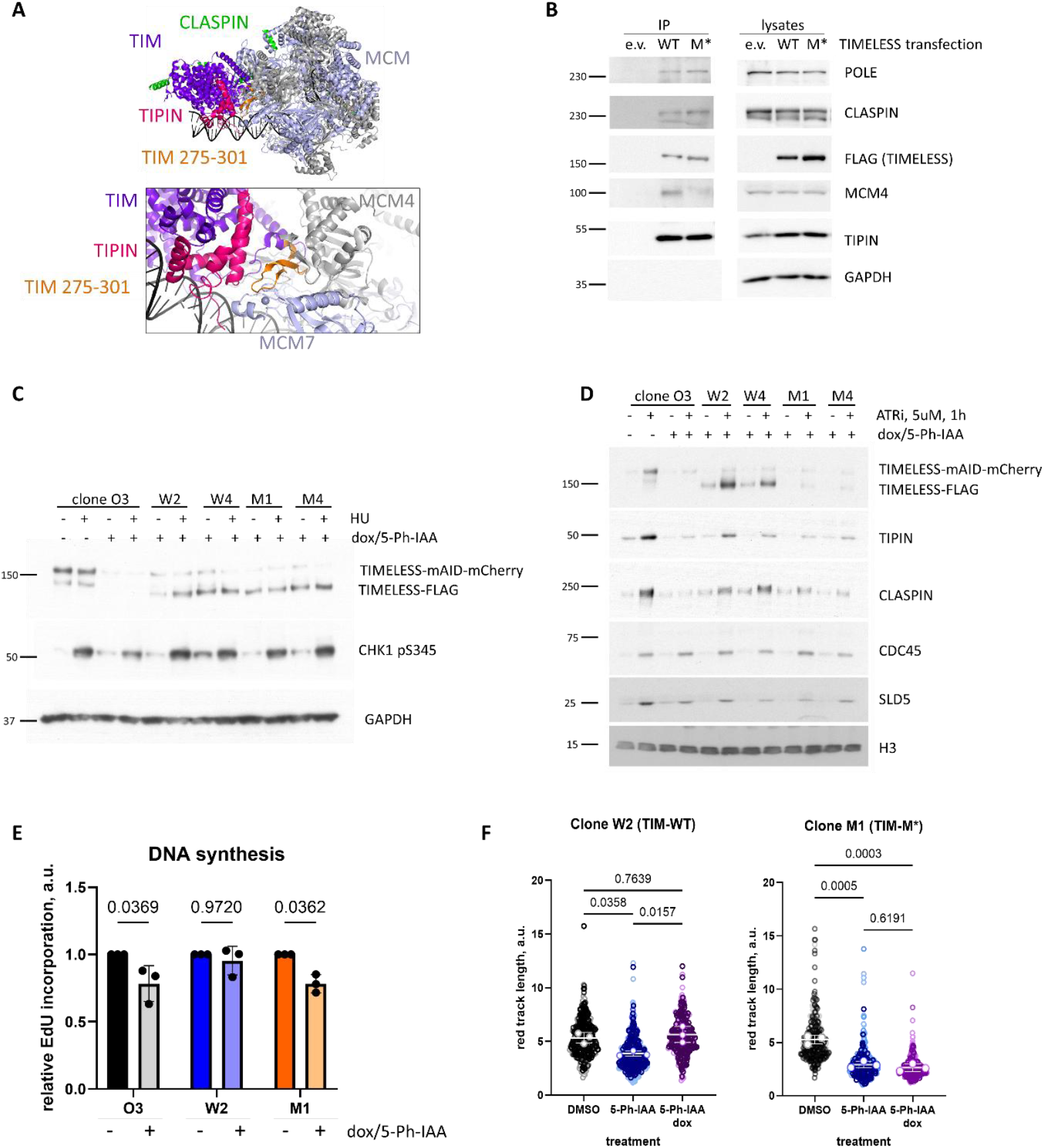
TIMELESS-MCM interaction is essential for regulation of replication fork speed, but not for replication checkpoint signaling. **A**. The position of the deletion in TIMELESS resulting in the disruption of TIMELESS-MCM interaction is shown on the experimental structure (PDB:7PFO (37)). **B**. Constructs expressing FLAG-tagged WT TIMELESS or TIMELESS-M* were transfected in 293T cells. 48h after transfection, TIMELESS was immunoprecipitated and eluted from the M2 beads using FLAG peptide. Western blot analyses of the indicated proteins are shown. **C-F**. mAID2 clone O3, and clones based on O3 expressing WT TIMELESS (W2 and W4) or TIMELESS M* (M1 and M4), were treated with 100 ng/ml doxycycline and 5Ph-IAA for 16h. 2mM HU was added for 1h as indicated. Western blot analyses on total cell lysates are shown **(C)**. ATRi or DMSO was added for 1h. Western blot analyses of nuclease-insoluble chromatin fractions are shown **(D)**. After a 30 min EdU pulse, quantification of EdU incorporation by FACS is shown, based on three independent experimental repeats **(E)**. Cells were pulsed with 20 µM CldU for 10 min followed by 200 µM IdU for 20 min, and lysed followed by DNA fiber analysis. Quantification of red track length (CldU) based on three independent experiments, is shown **(F)**. One-way ANOVA (**E**) or two-way ANOVA (**F**) were used for statistical analyses

To further test the effect of disrupting the interaction between TIMELESS and MCM on DNA replication, we expressed WT TIMELESS or TIMELESS-M* under the doxycycline-inducible promoter on the mAID2 clone O3 able to deplete endogenous TIMELESS. We initially selected two clones for each variant of TIMELESS – W2 and W4 expressed WT TIMELESS while M1 and M4 expressed TIMELESS-M*. We confirmed that these clones could efficiently degrade endogenous TIMELESS-mAID mCherry after 5-Ph-IAA treatment and express WT or M* TIMELESS after doxycycline treatment (**Fig. 5C**). The expression of FLAG-tagged dox-inducible TIMELESS was comparable to the expression of TIMELESS-mAID-mCherry before the depletion (**Fig. 5C**). While TIMELESS depletion in clone O3 decreased CHK1 phosphorylation after replication stress induction by hydroxyurea (HU) treatment, expression of either WT or M* TIMELESS was sufficient to support checkpoint signaling activation (**Fig. 5C**). This confirmed that the interaction between MCM and TIMELESS was not essential for the function of TIMELESS in the replication checkpoint.

Next, we wanted to check whether the interaction between MCM and TIMELESS is essential for FPC chromatin loading. We used ATRi-induced origin firing to observe the loading of replisome components onto chromatin. Disruption of MCM-TIMELESS interaction strongly decreased TIMELESS chromatin loading in response to ATRi, while recruitment of TIPIN and CLASPIN was somewhat decreased (**Fig. 5D, S5B**). Recruitment of CMG components CDC45 and SLD5 to chromatin, and an increase in EdU incorporation in response to ATRi were not affected by the disruption of TIMELESS-MCM interaction (**Fig. 5D, S5B**), in agreement with TIMELESS not being required for origin firing (**Fig. 2E-G, S2A-B**).

Just as TIMELESS-depleted cells (**Fig. S1B**), TIMELESS-M*-expressing cells showed a significant decrease in proliferation (**Fig. S5D**) and overall DNA synthesis (**Fig. S5E**), suggesting that the interaction between TIMELESS and MCM is essential for cell growth and DNA replication speed. We chose clones W2 and M1 for further analysis because they showed similar proliferation rates without treatments (**Fig. S5D**).

Since it has been shown before that TIMELESS depletion decreases replication fork speed, we decided to check whether TIMELESS-M* could support full replication fork speed. DNA fiber analysis confirmed that TIMELESS depletion decreased replication fork speed (**Fig. 5F**). While expression of WT TIMELESS fully restored replication fork speed, expression of TIMELESS-M* mimicked TIMELESS depletion, with significantly decreased replication fork speed (**Fig. 5F**). These data indicate that the interaction between MCM and TIMELESS is essential for supporting full replication fork speed in human cells.

## Discussion

### The effect of doxycycline on origin licensing and S-phase entry

Our data indicate that doxycycline treatment, and not TIMELESS depletion, led to a delay in licensing (**Fig. S1E-F**) and resulted in delayed DNA synthesis and S-phase entry in mAID1 clones (**Fig. 1D, E)**. Interestingly, delayed licensing did not affect CMG assembly as indicated by unperturbed CDC45 chromatin recruitment (**Fig. 2A-B**), but it did delay PCNA chromatin binding (**Fig. 2A-B**). Based on these data, in our experiments, in agreement with (41), replication complexes assemble before licensing is completed, but they are inhibited to prevent re-replication (41). Our data indicate that PCNA is only recruited to chromatin after DNA synthesis is initiated, which is delayed by doxycycline treatment in our experiments. Moving forward, we primarily use the mAID2 system, and lower doxycycline concentrations to 100ng/ml, where possible. The effect of doxycycline on origin licensing is an important consideration for all cell cycle and DNA replication researchers using TET-inducible systems.

### Chromatin loading of the FPC during the initiation of DNA replication

The roles of TIMELESS in replication fork progression include overcoming intrinsic barriers such as secondary structures and DNA-protein complexes (39), which could cause fork deceleration. However, biochemical experiments (9) demonstrated that TIMELESS is a key component in the FPC that maintains full fork speed even in the absence of obstacles. We therefore wanted to check whether the decrease in DNA synthesis that we observed was due to a failure to assemble a functional FPC at the replication fork. Our data showed that TIMELESS depletion led to a depletion of its partner TIPIN, and a decrease in chromatin binding of CLASPIN both during S-phase entry and during ATRi-induced replication origin firing (**Fig. 2**). TIPIN has been shown to recruit CLASPIN to ssDNA during replication checkpoint activation (4); however, the roles of TIMELESS/TIPIN in CLASPIN loading during the initiation of DNA replication have not been previously described. Interestingly, previous studies demonstrated that CLASPIN, loss does not affect TIMELESS-TIPIN loading (5), confirming that the TIMELESS-TIPIN complex, and not CLASPIN mediated the FPC chromatin association. Our systems do not allow us to determine whether it is TIMELESS or TIPIN that is responsible for CLASPIN chromatin loading. Despite trying to disrupt all known contacts between TIMELESS and CLASPIN, we have not been able to create a mutant version of TIMELESS unable to bind CLASPIN. Perhaps, future structural studies may guide this direction of research more efficiently. Our data indicate that the presence of TIMELESS/TIPIN is critical for CLASPIN chromatin loading during replication initiation to ensure full replication fork speed. Our study was also able to expand this observation to the second FPC instance, as low levels of TIMELESS-M* chromatin binding supported similarly low levels of CLASPIN and TIPIN chromatin association (**Fig. 5**). Our data support the model where the role of TIMELESS/TIPIN in checkpoint regulation could be through recruiting CLASPIN to ssDNA/RPA.

Synchronization experiments observing chromatin loading of the FPC components during S-phase entry showed that the FPC was present on chromatin before the loading of DNA polymerase alpha – the enzyme responsible for the initial DNA synthesis on both strands during replication initiation (**Fig. 3A-B**). Together with the FPC loading requiring CDK2 or CDC7/CDK1 kinase activities (**Fig. 3C-F**), this suggested a model where FPC is loaded after CMG assembly/activation but before DNA polymerase alpha. However, inhibition of DNA synthesis by aphidicolin strongly decreased FPC chromatin loading (**Fig. 3C-F**), indicating that DNA synthesis is required for full FPC association with chromatin. This stepwise loading of the FPC is consistent with the presence of additional FPC instances on the ssDNA between Okazaki fragments. The leading edge FPC would load at the time of the CMG activation, as soon as the leading edge is established, and the ssDNA-associated FPC would load after DNA synthesis starts creating ssDNA spaces between Okazaki fragments. In agreement with this model, loading of both FPC instances would require CMG activation and kinase activities of CDK2 and CDC7/CDK1, while the second FPC would only be recruited once DNA synthesis starts and would therefore be blocked by aphidicolin.

### Position of TIMELESS at the replication fork

TIMELESS, as a component of the FPC, has been well described for its role in fork stabilization, interaction with DNA polymerases, as well as checkpoint activation (reviewed in (39)). These roles are not fully in agreement with its position at the leading edge of the replication fork, where it has been repeatedly identified by highly reliable structural studies (23–25). Additionally, TIPIN, TIMELESS’ obligatory binding partner, has been demonstrated to bind both dsDNA at the leading edge of the replication fork (23–25), and ssDNA/RPA (4), which is difficult to imagine simultaneously. Due to these inconsistencies, as well as our observation of a stepwise chromatin loading of TIMELESS during replication initiation, we used split-TurboID to shine more light on the position of TIMELESS at the replication fork. Surprisingly, the N-terminus of TIMELESS appeared to be in proximity to the C-terminus of AND-1, forming a functional biotin ligase that labeled replicative DNA polymerases, CDC45, CHK1, and other replication proteins (**Fig. 4**).

AND-1 is sometimes regarded as one of the FPC components (41) despite there being no structural evidence of direct interactions between AND-1 and TIMELES-TIPIN-CLASPIN. CLASPIN co-purified with AND-1 in co-immunoprecipitation experiments (42), which appeared to pull down the full replisome. A study in Xenopus egg extracts was able to co-purify AND-1 and TIPIN (40). Since both AND-1 and TIMELESS-TIPIN-CLASPIN interact with the CMG helicase, these co-purifications do not necessarily prove a direct interaction. One study indicated that AND-1 and CLASPIN both accumulate on ssDNA to promote S-phase checkpoint activation (43), which suggests a functional association, but on ssDNA and not at the leading edge of the fork where the FPC interacts with dsDNA (23–25). AND-1 could interact with ssDNA in its canonical position, but the reported accumulation of AND-1 on ssDNA (43) suggests that perhaps additional monomers or trimers of AND-1 are recruited to ssDNA during replication stress. Our successful combination of AND-1-TurboC/TurboN-TIMELESS in the absence of replication stress suggested that AND-1- and TIMELESS co-localized at unperturbed replisomes. While we cannot make a definitive conclusion, it is possible that the N-terminus of the leading-edge TIMELESS was in proximity to the long and flexible C-terminus of AND-1. However, POLD was biotinylated by the TIMELESS/TIMELESS combination, but not the TIMELESS/AND-1 combination, which indicates that the TIMELESS/TIMELESS contact was close to the lagging strand, while the TIMELESS/AND-1 and the TIMELESS/ORC6 contacts were not. We considered a possibility that two TIMELESS molecules coming together to produce this biotinylation could be from two different replication forks; however, if both tagged molecules of TIMELESS were located at the leading edges of their respective forks, biotinylation of DNA polymerase epsilon and CDC45, located on the directly opposite side of the CMG, is still very difficult to explain. Given that TIMELESS tagged with a shorter, N-terminal part of the TurboID, at its N terminus, was used in all three combinations, this strongly supports our model where there are two instances of TIMELESS/TIPIN at the replication fork. We propose that there is an additional TIMELESS-TIPIN-CLASPIN complex at ssDNA/RPA on the lagging strand, mediated by TIPIN-RPA interaction and supporting the background level of CHK1 phosphorylation by ATR in the absence of replication stress, known to be essential to suppress excessive origin firing (35).

### Significance of TIMELESS-MCM interaction

Previous studies describing TIMELESS depletion phenotypes (5, 37, 38, 44–46) documented the effect of depletion of both leading edge TIMELESS as well as lagging strand TIMELESS using either siRNA or a degron strategy, without separating the distinct roles of the two instances. Our system allowed us to specifically disrupt the leading edge TIMELESS without affecting the RPA-associated FPC. Our data indicated that leading edge TIMELESS is necessary for maintaining full speed of the replication forks, as well as efficient chromatin loading of TIPIN and CLASPIN. However, TIMELESS-M*, unable to interact with MCM, was sufficient to support the checkpoint function of TIMELESS, likely mediated by CLASPIN loading at ssDNA to support CHK1 phosphorylation by ATR (43). Future studies will uncover the effect of disrupting TIPIN-RPA interaction that should block the checkpoint function of the FPC while preserving the fork speed regulated by the leading edge FPC (this has been shown biochemically in (4)) – our TIMELESS-centered system does not allow us to easily check that in cells. Nevertheless, we show here for the first time that the checkpoint function of the FPC does not require its interaction with the CMG and is mediated by a separate instance of TIMELESS. Given that TIMELESS is frequently overexpressed in cancers, our study has important implications for how the excess of TIMELESS protects cancer cells from DNA damage – without changing the number of active replication forks, additional FPC complexes at each fork could strengthen the checkpoint signaling, providing resistance to DNA damage, including chemotherapy.

## Contributions

S.V., K.S., R.H., N.R.C., S.S.M., A.M., Si.V., and T.N.M. performed experiments and analyzed data. S.V. designed experiments and wrote the initial draft of the manuscript. T.N.M. conceived and directed the project, acquired funding, designed experiments, reviewed and edited the manuscript.

## Funding

This work was supported by Estonian Research Council (research grant PRG1477 to T.N.M) and startup funding from UPMC Hillman Cancer Center to T.N.M. Research reported in this publication was supported by the National Cancer Institute of the National Institutes of Health under Award Number P30CA047904.

## Data availability statement

The data underlying this article are available in the article and in its online supplementary material.

## Notes

### Competing Interest Statement

The authors have declared no competing interest.

