## Supplementary figures for "A new model for coordinating the functions of TIMELESS at the replication fork"

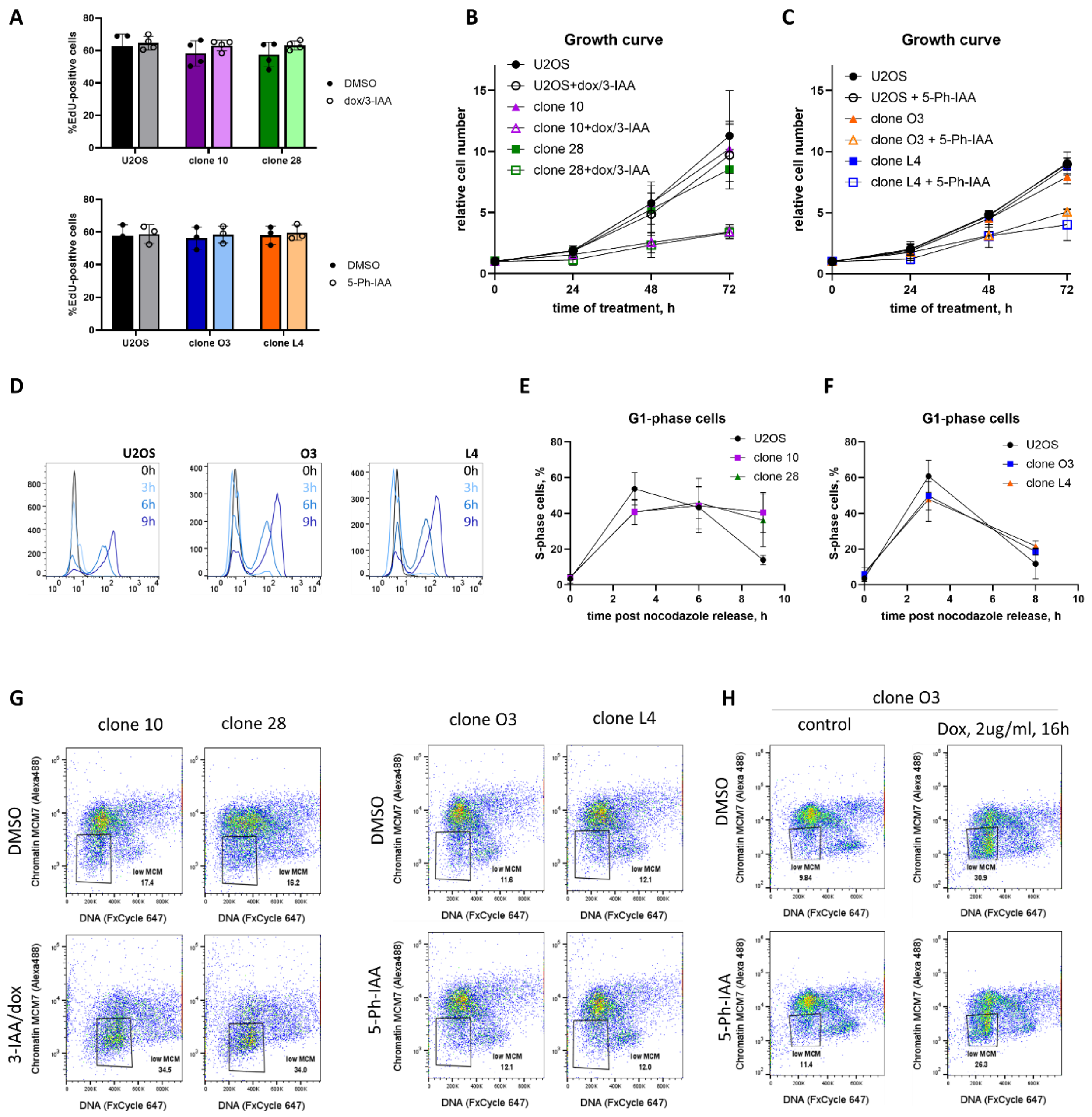


**Figure S1. A.** U2OS, mAID1 clones 10 and 28, or mAID2 clones O3 and L4 were treated for 16 h with dox/ 3-IAA, auxin 5-pH-IAA. 10 µM EdU was added for the last 30 min of treatment. Quantification of the flow cytometry data is shown- mean + SD from n = 3 independent experiments. **B-C.** Equal numbers U2OS, mAID1 clones 10 and 28, or mAID2 clones O3 and L4 were seeded on 60 mm dishes and treated with DMSO or dox/3-IAA (mAID1) or 5-Ph-IAA (mAID2) for 72 h. Cell numbers were counted every 24 h and growth curves were plotted. The data are depicted as mean + SD from n = 3 independent experiments. **D-F.** U2OS, mAID1 clones 10 and 28 or mAID2 clones O3 and L4 were synchronized using the double thymidine/nocodazole block, and treated as indicated on **Fig.1C**. 10 µM EdU was added for the last 30 min of treatment. Quantification of the flow cytometry data is shown - mean + SD from n = 3 independent experiments **(E-F).** Cells were assigned to G1-phase based on DNA content of 2n and being EdU negative. EdU incorporation histograms are shown for synchronized untreated U2OS and clones O3 and L4 showing that TIM tagging did not affect the S-phase entry without TIMELESS depletion **(D)**. **G-H.** mAID1 clones 10 and 28, or mAID2 clones O3 and L4 were treated for 16 h, as indicated. Cells were extracted, MCM7 was stained with antibodies, DNA was stained with FxCycle FarRed dye. Presence of MCM7 on chromatin was analyzed by FACS.

**A**

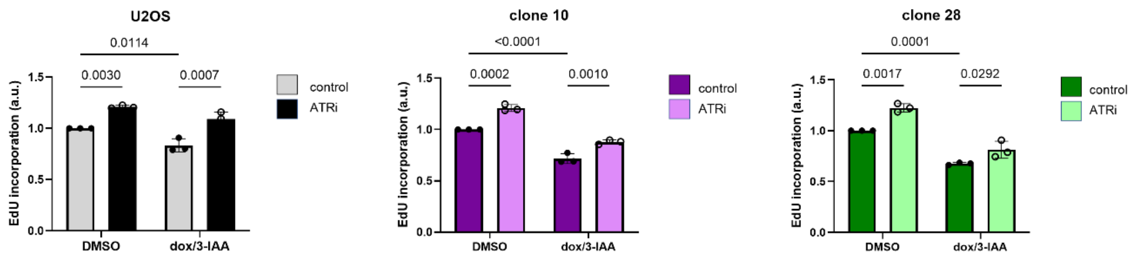


**B**

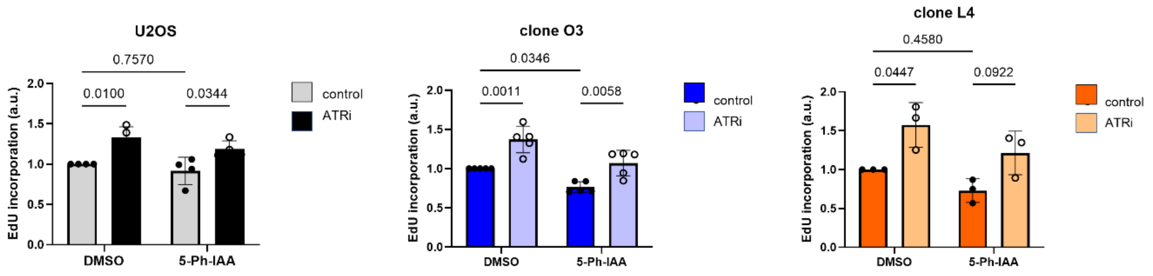


**Figure S2. A-B**. U2OS, mAID1 clones 10 and 28 **(A),** or mAID2 clones O3 and L4 **(B)** were treated for 16h with dox/3-IAA, or 5-Ph-IAA, as indicated. 5 µM ATRi was added to the indicated samples for 1 h, and 10 µM EdU was added for the last 30 min of treatment. Flow cytometry plots showing relative EdU incorporation are shown. Mean + SD from n = 3 - 5 independent experiments are shown.

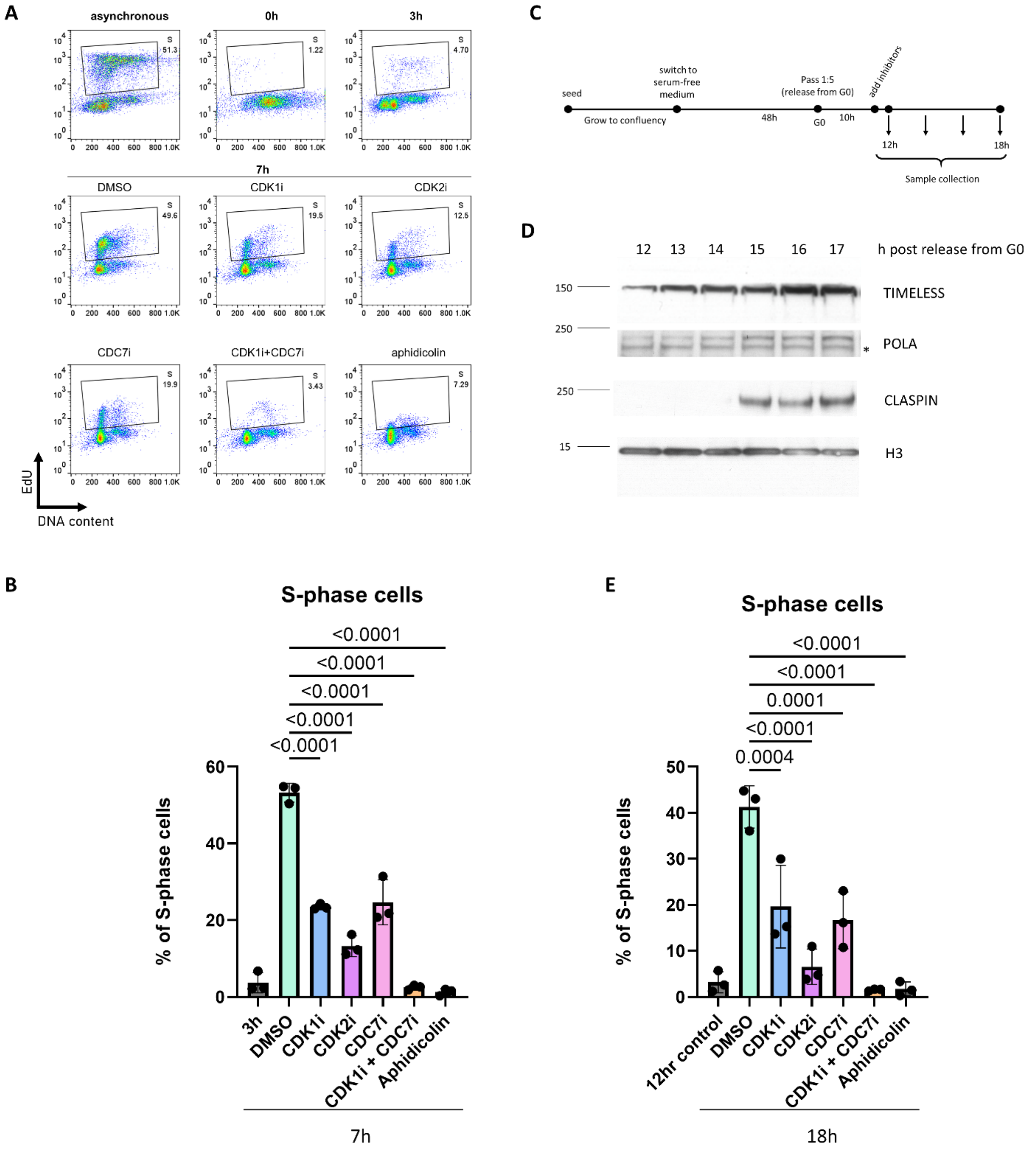


**Figure S3. A-B**. U2OS cells were synchronized as indicated on **Fig.1C.** 3h post nocodazole release, the indicated inhibitors were added. Samples were collected at the indicated timepoints post nocodazole release. 10 μM EdU was added for the last 30 min of treatment. Plots (**A**) and quantification (**B**) of the flow cytometry data is shown - mean + SD from n = 3 independent experiments. **C.** RPE hTERT synchronization strategy. **D-E.** RPE-hTERT cells were synchronized by contact inhibition and serum starvation as indicated on **Fig S3C.** Cells were collected at indicated timepoints, chromatin fraction was isolated and analyzed by western blotting **(D).** Indicated inhibitors were added 10h after release from G0, samples were collected at indicated timepoints after 30 min pulse of EdU. Quantification of the flow cytometry data is shown - mean + SD from n = 3 independent experiments **(E).**

**Figure S4: A-B.** HEK293FT cells were transfected with the plasmids expressing indicated TurboN- or TurboC-tagged proteins. 48h after transfection cells were treated with 50 μM biotin for 1 hour and lysed. Cell lysates were analyzed by western blot using antibodies against indicated proteins.

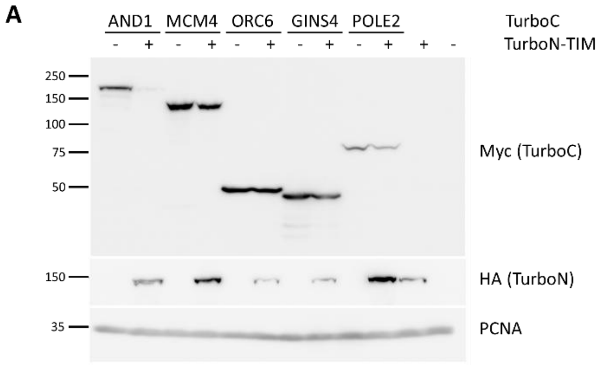


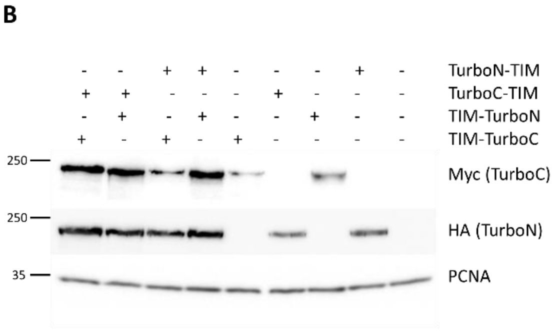

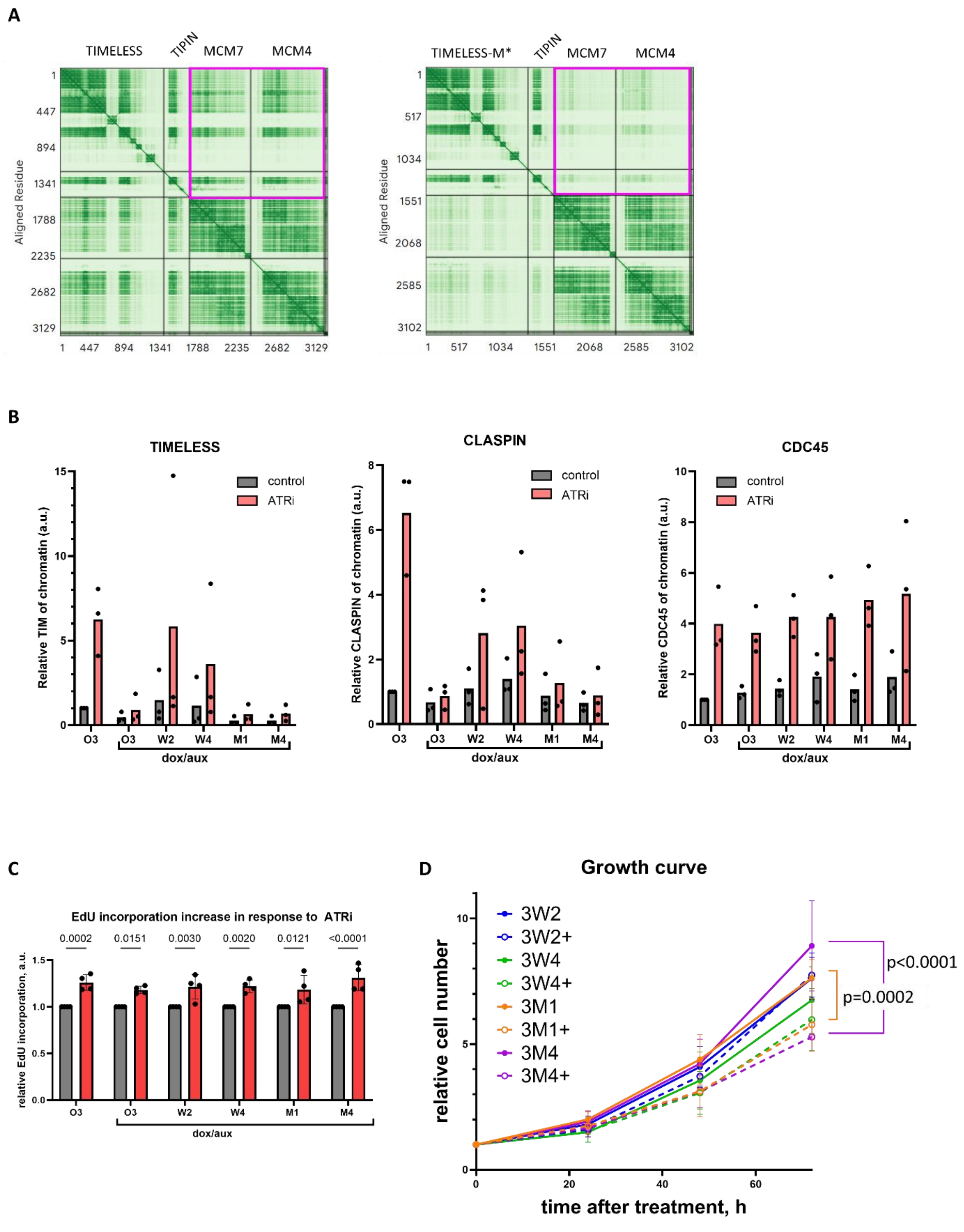


**Figure S5. A.** TIPIN, MCM7 and MCM4 are modeled in Alphafold3 with TIMELESS WT or TIMELESS-M*. The intensity of green indicated the predicted error. Magenta frame indicates the relative position confidence between TIMELESS and MCM subunits. **B.** Quantification of western blots from **Fig. 5D**. Data for three independent experiments are shown. **C.** mAID2 clone O3, and clones based on O3 expressing WT TIMELESS (W2 and W4) or TIMELESS M* (M1 and M4), were treated with doxycycline and 5Ph-IAA for 16h, followed by the incubation with DMSO or ATRi for 1h. After a 30 min EdU pulse, quantification of relative EdU incorporation by FACS is shown, based on three independent experimental repeats. **D.** Equal numbers mAID2 clone O3, and clones based on O3 expressing WT TIMELESS (W2 and W4) or TIMELESS M* (M1 and M4), were seeded on 60 mm dishes and treated with DMSO or dox/5-Ph-IAA for 72 h. Cell numbers were counted every 24 h and growth curves were plotted. The data are depicted as mean + SD from n = 3 independent experiments.
